# A free boundary model of epithelial dynamics

**DOI:** 10.1101/433813

**Authors:** Ruth E Baker, Andrew Parker, Matthew J Simpson

## Abstract

In this work we analyse a one-dimensional, cell-based model of an epithelial sheet. In this model, cells interact with their nearest neighbouring cells and move deterministically. Cells also proliferate stochastically, with the rate of proliferation specified as a function of the cell length. This mechanical model of cell dynamics gives rise to a free boundary problem. We construct a corresponding continuum-limit description where the variables in the continuum limit description are expanded in powers of the small parameter 1/*N*, where *N* is the number of cells in the population. By carefully constructing the continuum limit description we obtain a free boundary partial differential equation description governing the density of the cells within the evolving domain, as well as a free boundary condition that governs the evolution of the domain. We show that care must be taken to arrive at a free boundary condition that conserves mass. By comparing averaged realisations of the cell-based model with the numerical solution of the free boundary partial differential equation, we show that the new mass-conserving boundary condition enables the coarsegrained partial differential equation model to provide very accurate predictions of the behaviour of the cell-based model, including both evolution of the cell density, and the position of the free boundary, across a range of interaction potentials and proliferation functions in the cell based model.

## 1. Introduction

Cell biology experiments typically produce complex, quantitative experimental data that can include both cellular-level and tissue-level information [8, 11, 12, 23]. However, it can often difficult to integrate these multi-scale data to give new insights. This challenge provides a clear motivation for the use of mathematical models where individual, cell-based mechanisms can be implemented and explored in a computational framework [1, 17, 21, 24]. This approach can allow us to qualitatively explore the relationship between individual-level properties and population-level outcomes using repeated computational simulations as well as comparing predictions of different models that act at different scales [14, 18]. Furthermore, it is possible to provide a quantitative, more rigorous mathematical connection between the individual-level properties and population-level outcomes by using coarse-graining techniques to derive an approximate continuum-limit description of the individual-level description [13, 15, 16].

Depending on the biological context, there are many different kinds of individual-based models that can be used to simulate cell biology processes including random walk frameworks involving point particles [2, 3] or random walk frameworks based on an exclusion process that explicitly account for excluded volume effects as well as the shape and size of the individuals in the system [9, 19]. While discrete models based on point particles and exclusion processes have been successfully applied to study many cell biology phenomena, these models do not include any mechanical effects that are known to be important in a host of applications. For example, tissue stiffness is known to play a key role in epithelial cancer progression, with different rates of invasion associated with different tissue stiffness conditions [20]. Cancer detection is another clinical application where tissue mechanics and tissue stiffness, in terms of mammographic density, is thought to be associated with breast cancer risk [7]. Therefore, for certain applications, it is relevant to use a mechanical framework to study the motion and interaction of individual cells rather than focusing on a random walk framework.

In this work we re-examine a mechanical model of epithelial tissue mechanics first presented by Murray [13, 15]. The model describes a one-dimensional population of cells, where nearest neighbour cells interact through a force potential and the motion of each individual is governed by an overdamped, deterministic equation of motion. Like Murray [13, 15], we consider the case where the left-most boundary of the population of cells is fixed at *x* = 0, and the right-most boundary is a free boundary, *x* = *L*(*τ*), where *τ* is time. In the first part of our work we consider a non-proliferative population where individual cells undergo movement only. In this context the cell-based model is deterministic and the evolution of the free boundary is the net result of the deterministic interactions between the *N* individuals. In the second part of our work we consider a population of cells that is both proliferative and motile, and in this context the individual cell based model is stochastic. Here the evolution of the cell density and the position of the free boundary is the net result of a combination of the deterministic motility mechanism and stochastic proliferation events, where the rate of proliferation is taken to be a function of the length of each cell in the stochastic simulations. In all cases considered we study expanding populations where *L*(*τ*) is an increasing function of time.

The key focus of this work is the derivation of a continuum-limit partial differential equations (PDE) description of the individual-based model that provides an accurate description of both the macroscopic density of cells within the domain, as well the movement of the free boundary, *L*(*τ*). We make progress by defining continuous functions by expanding in powers of the small parameter, 1/*N*, so that, formally, our continuum limit description is accurate in the limit, *N* → ∞ [5]. By carefully neglecting terms of 𝒪(1/*N*^2^), we derive a free boundary problem that describes the spatial and temporal evolution of the cell density within the domain, 0 < *x* < *L*(*τ*), as well as the temporal evolution of the free boundary, *L*(*τ*). We show that our new free boundary condition conserves mass. Comparing averaged data from cell-based simulations with the numerical solution of the continuum limit PDE description of the free boundary problem confirms that the new mass-conserving boundary condition provides an accurate description of the dynamics of the cell-based model across a range of different individual-based mechanisms conditions.

## 2. A discrete model of cell dynamics in one dimension

In this work, we consider one of the simplest cell-based, off-lattice models of a onedimensional epithelial cell population that captures cell-cell adhesion interactions and bulk cellular elasticity. Cells occupy volume and can undergo deformation, neighbouring cells come into contact with each other at node points (Figure 1), and they interact with each other and the local microenvironment [13]. We formulate a mathematical description of the dynamics of the cell population using *x*_*i*_ to denote the position of node *i* (Figure 1).

From Newton’s second law of motion

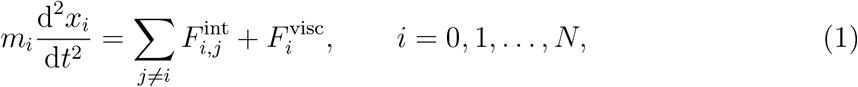

where 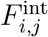, the force node *j* exerts on node *i*, represents the combined effects of cellular bulk elasticity and cell-cell adhesion^1^, 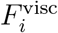 is the viscous force acting on the *i*^th^ node, and *m*_*i*_ is the mass associated with the *i*^th^ node.

**Figure 1:**
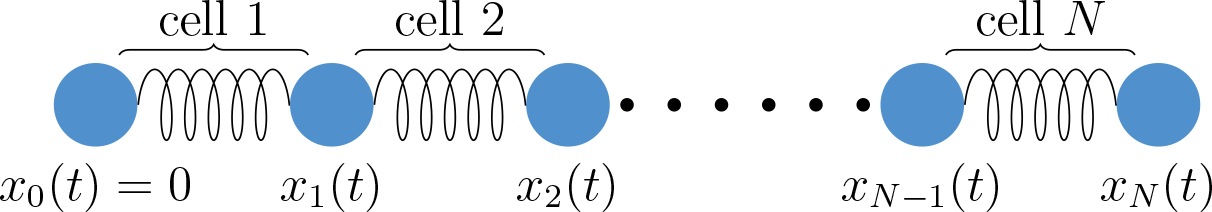
Schematic of the cell-based model where the cells are here represented by springs and nodes the points where two cells touch. There are *N* cells, and node positions are denoted by *x*_*i*_, for *i* = 0,1,…,*N*, with the left boundary of the first cell (*i.e.* node 0) fixed at the origin so that *x*_0_(*t*) = 0.

We now make a number of further assumptions to simplify equation (1). Firstly, we assume that cells interact with only their nearest neighbours, so that 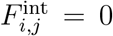 for *j* ≠ *i* ± 1, that cells cannot exchange neighbours, and that node zero is pinned at the origin. This entails 0 = *x*_0_(*t*) < *x*_1_(*t*) < … < *x*_*N*−1_(*t*) < *x*_*N*_(*t*). Secondly, we assume that the viscous force, 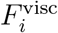, which is generated by cell-cell and cell-matrix interactions, can be modelled as proportional to velocity, d*x*_*i*_/d*t*, with viscosity coefficient *η*. Thirdly, cells move in dissipative environments, so we assume *m*_*i*_d^2^*x*_*i*_/d*t*^2^ ≈ 0. Finally, we assume the cell population is homogeneous, so that *m*_*i*_ = *m* for *i* = 0,1,…, *N*, and cells respond and generate forces according to the same physical law. As a result, the dynamics of the population can be modelled using the following system of ODEs:

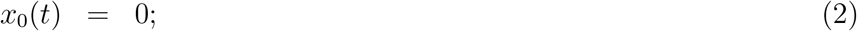

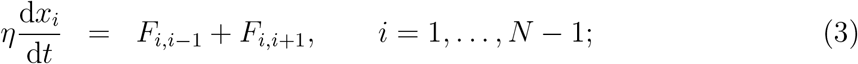

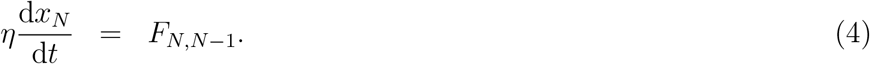

Note that we have suppressed the superscript in 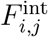 for clarity from this point onwards. The system is closed by specifying appropriate initial conditions, 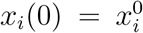 for *i* = 1, …, *N*. To provide a simple exposition, in the initial stages of this work we will assume that cells can be modelled as linear springs; extension to more general cases is provided in Section 4.

## 3. Linear force law model

We first assume that the interaction force between cells *i* and *i* ± 1 can be modelled using a linear force law with constant *k* > 0 and equilibrium length *a* > 0, as in [13], so that

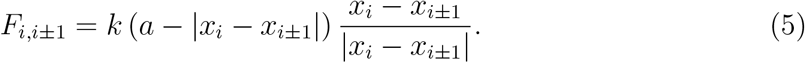

Letting *α* = *k*/*η* we have

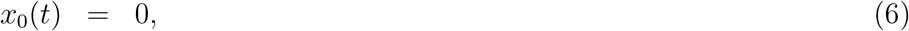

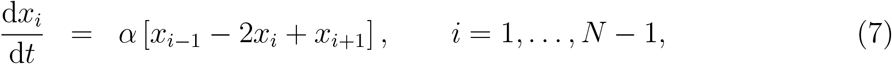

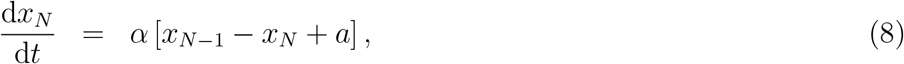

with initial conditions, 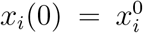 for *i* = 1,…, *N*.

System (6)-(8) can be solved analytically. However, since in this work our aim is to extend to more general (analytically intractable) cases where nonlinear force terms are used to model cellular dynamics, we solve for the position of each node, *x*_*i*_, numerically using a simple forward Euler method with time-step Δ*t* = 0.001. Exemplar results for the model are shown in Figure 2, where we demonstrate how the leading edge and cell density of an initially compressed population of cells evolves over time.

### 3.1. Continuum approximation

To make progress in deriving an equivalent continuum, coarse-grained model, with a slight abuse of notation we will extend node position, *x*_*i*_(*t*), which is only defined for discrete *i* ∈ {0,…, *N*}, to a smooth function, *x*(*i*, *t*), which is defined for *i* ∈ [0, *N*]^2^. The function *x*(*i*, *t*) will approximate *x*_*i*_ when *i* is an integer:

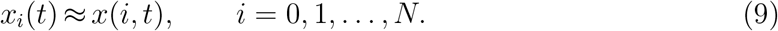

To facilitate coarse graining, we first non-dimensionalise the model specified in equation (6)-(8) using the scalings

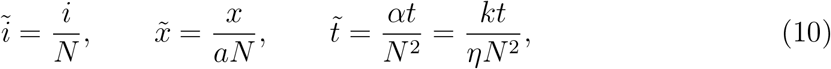

so that 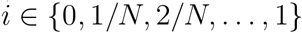 and 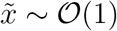. The model equations are then

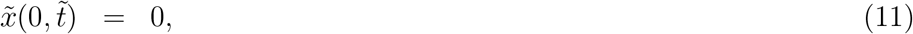

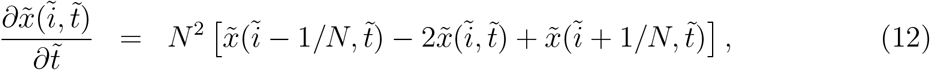

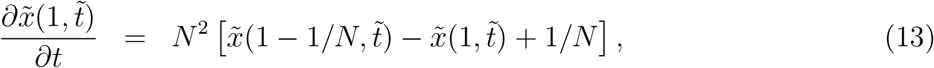

where equation (12) holds for 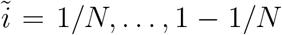, and we have corresponding initial conditions of the form 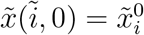 for *i* = 1/*N*,…, 1.

Performing a Taylor expansion about 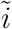 within equation (7) gives, on neglecting terms that are 𝒪 (1/*N*^2^),

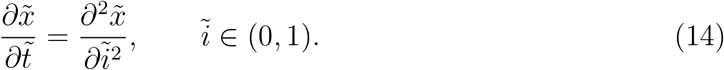

The left-hand boundary condition is simply 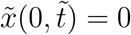. To derive the right-hand boundary condition, we again Taylor expand and neglect terms that are 𝒪 (1/*N*^2^) to give, at 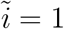,

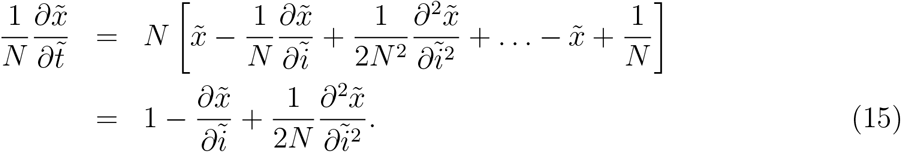

In terms of the dimensional variables, the coarse-grained model is therefore

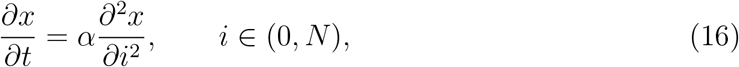

with boundary conditions

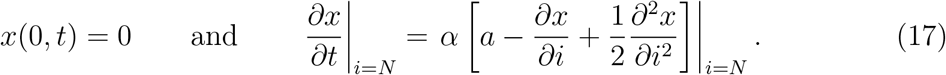

Initial conditions can be specified by extending the discrete initial conditions, 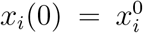 for *i* = 1,…, *N*, to a continuous function *x*(*i*, 0) such that 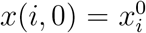 for *i* = 1,…, *N*.

Throughout this work, for simplicity we extend the discrete initial condition to a piecewise linear continuous function.

### 3.2. Derivation of the corresponding cell density model

Cell density can be defined implicitly using the relation

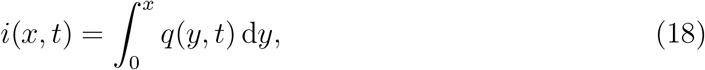

where *i* is the cell index. Equation (18) is equivalent to *q*(*x*, *t*) = ∂*i*(*x*, *t*)/∂*x*, and it ensures *i* = 0 at the left-hand boundary, *x* = 0. To reformulate equations (16) and (17) in terms of variation in cell density with position, *x*, and time, *t*, we follow [13] and perform a change of variables from (*i*, *t*) to (*x*, *τ*) where *i* and *x* are related through equation (18) and *t* = *τ*.

Noting that

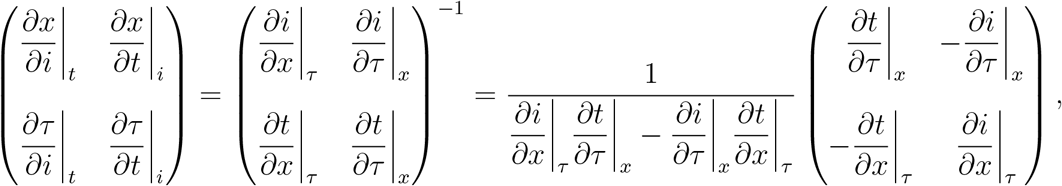

we have

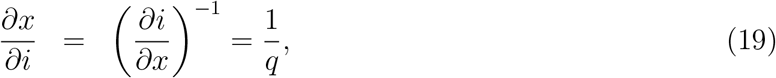

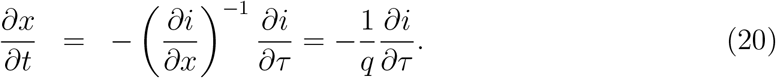

Substituting equation (19)-(20) into the right-hand side of equation (16) gives

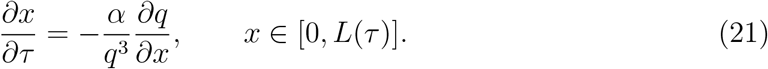

Equation (21) is the characteristic equation and it represents how the domain evolves over time through tracking constant node index, *i*. After a simple rearrangement (multiplying by *q* and differentiating with respect to *x*) we have

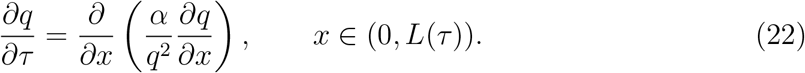

The same change of variables applied to the boundary conditions in equation (17) yields

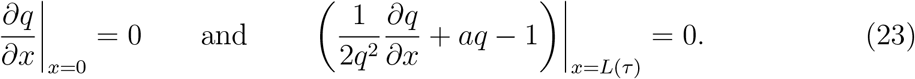

As a check on the validity of the derived boundary conditions, we note that the system must conserve total cell density, *i.e.*

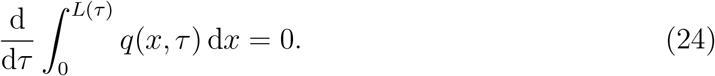

Evaluating the above expression gives (again, using the characteristic equation (21)

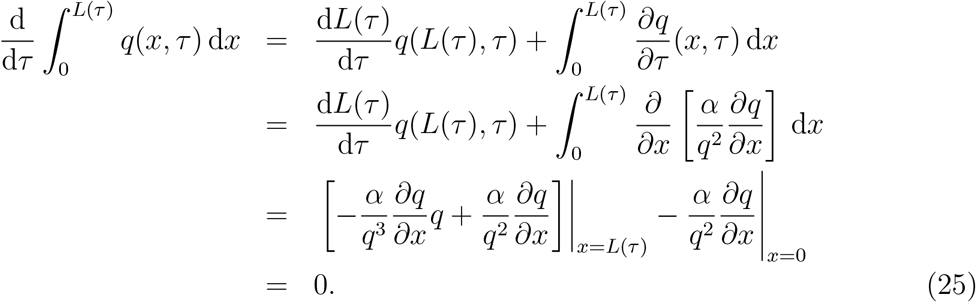

Therefore total density is conserved using the derived boundary conditions. Note that the boundary condition applied at the free, right-hand boundary, *L*(*τ*), is slightly different to that derived in [13], where the boundary condition was derived by neglecting terms that are 𝒪(1/*N*) and is of the form *q* = 1/*a* for *x* = *L*(*τ*).

To establish initial conditions, we use equation (18) together with a finite difference approximation to write

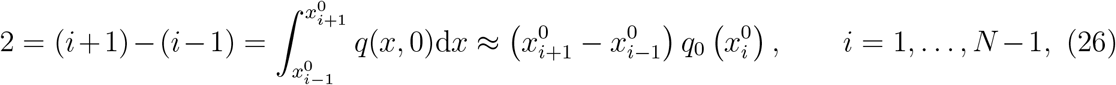

which can be rearranged to give

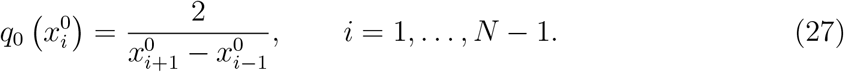

A similar finite difference approximation applied at the left- and right-hand boundaries gives

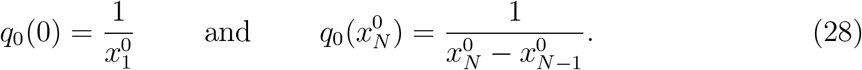

We treat *q*_0_(*x*) as piecewise linear between node positions.

#### 3.2.1. Numerical solution

In summary, the coarse-grained model consists of a PDE for the evolution of cell density

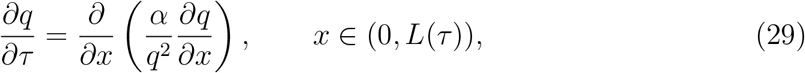

together with boundary conditions

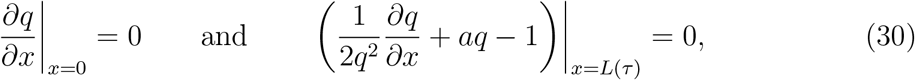

and initial condition

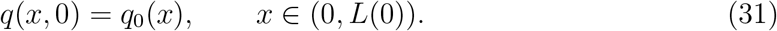

The characteristic equation is

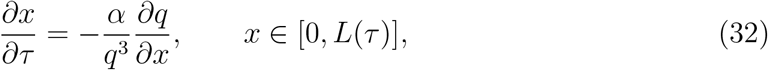

and we can use it to specify the evolution of the domain with time. In particular, we have

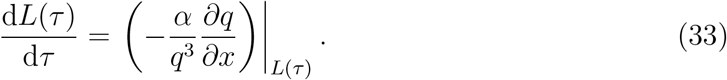

In order to solve equations (29)-(33) numerically we employ a Lagrangian transformation to map the free boundary problem to a fixed domain. We let *τ* = *T* and

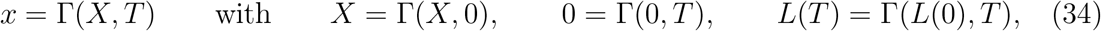

so that

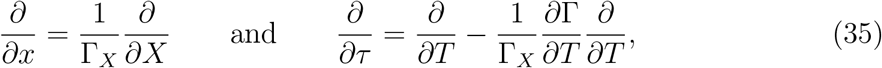

where we have adopted the notation ∂Γ/∂*X* = Γ_*X*_. Substitution into equations equations (29) and (33) yields equations for evolution of the domain and the density therein:

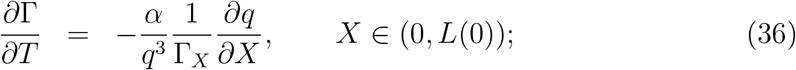

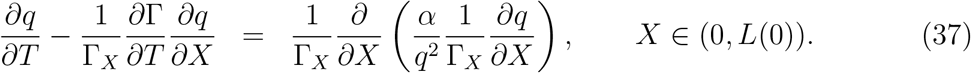

The initial and boundary conditions for Γ(*X*, *T*) are specified in equation (34), and for *q*(*X*, *T*) we have

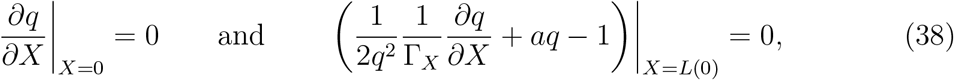

together with

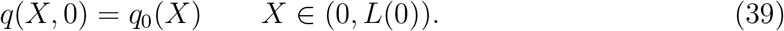

We solve the model numerically using an implicit finite difference method with Picard iteration. Full details are given in Appendix A.

### 3.3. Results

The coarse-grained PDE model is very accurate in its prediction of both evolution of the cell density, *q*(*x*, *τ*), and the free boundary at *x* = *L*(*τ*) (see Figure 2), even for relatively low cell numbers (here we show results for cell numbers as low as *N* = 15). The accuracy of the PDE model increases as the cell number, *N*, increases; this is in line with expectations since the error of the coarse-grained PDE model is 𝒪(1/*N*^2^). To ensure sensible comparisons, the results in Figure 2 were generated by initialising *N* cells with equal lengths in the interval *x* ∈ (0, 30) and varying the model parameters such that *α* = 15(*N*/45)^2^ and *a* = 45/*N*. This choice ensures that the scalings for *x* and *τ* do not change with increasing *N*. In each case, cells are initially compressed but will eventually expand to fill the domain *x* ∈ (0, 45).

We also compare the results of our model against those derived in [13], where the boundary condition at the free boundary was derived by neglecting terms 𝒪(1/*N*) and is of the form *q* = 1/*a* for *x* = *L*(*τ*). As expected, the boundary condition derived here leads to a more accurate prediction of the dynamics of the cell-based model because we neglect only terms that are 𝒪(1/*N*^2^) rather than 𝒪(1/*N*).

**Figure 2:**
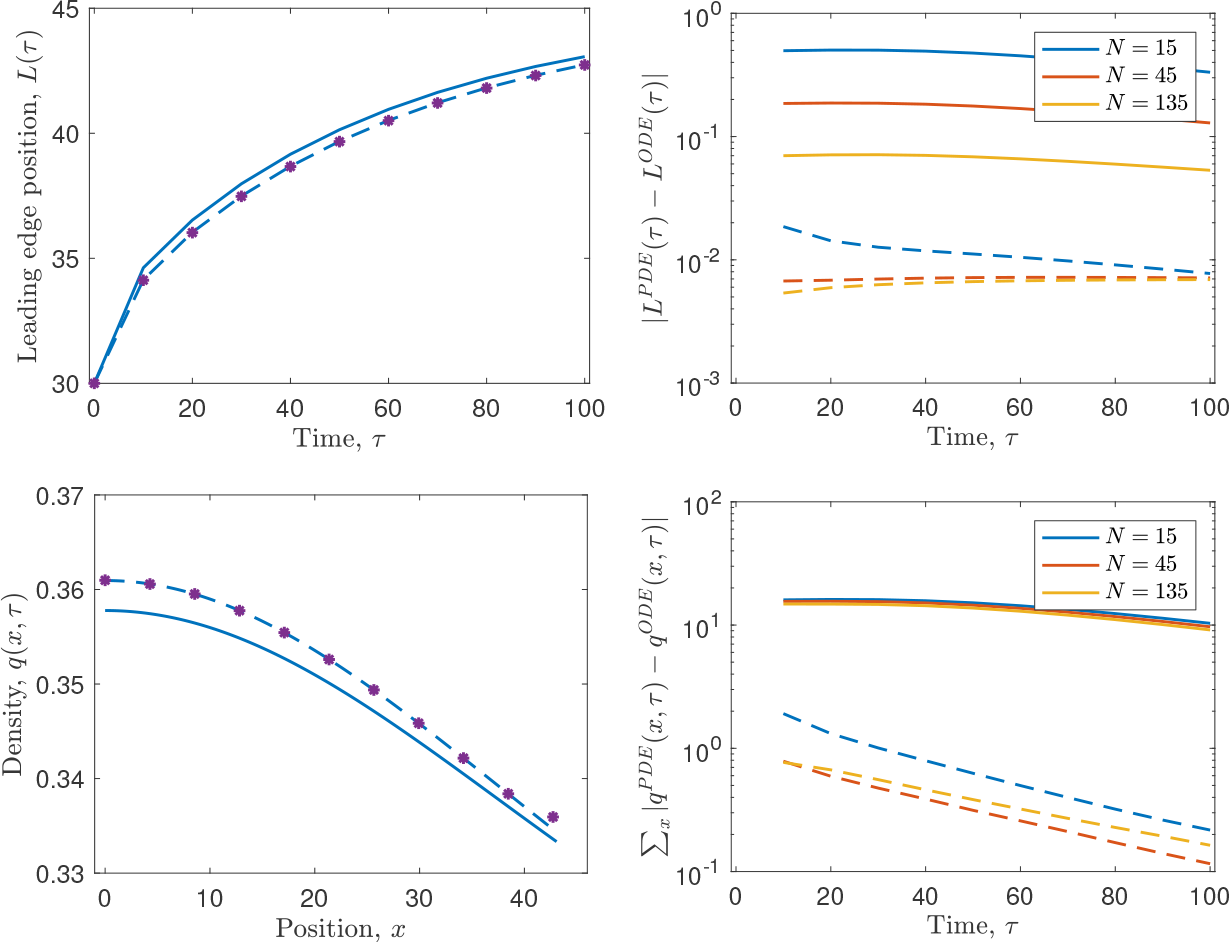
Comparison of the leading edge position, *L*(*τ*), and cell density, *q*(*x*, *τ*), predicted by the cell-based model, equation (6)-(8), and the coarse-grained PDE model, equation (29)-(33), as the cell number, *N*, is varied. In each case, *aN* = 45 and *α/N*^2^ = 135 are kept constant. On the left-hand side, the leading edge position, *L*(*τ*), predicted by the cell-based model with *N* =15 is plotted using purple asterisks, whilst the prediction of the PDE model using the boundary conditions derived in Section 3.1 and stated in equation (23) is plotted as a dashed blue line. For comparison, the leading edge position predicted using the boundary conditions of [13] is plotted as a solid blue line. On the right-hand side, the error in the predictions of the coarse-grained model are shown for both the boundary conditions stated in equation (23) (dashed lines), and those derived by [13] (solid lines), for a range of values of cell number, *N*.

**Figure 3:**
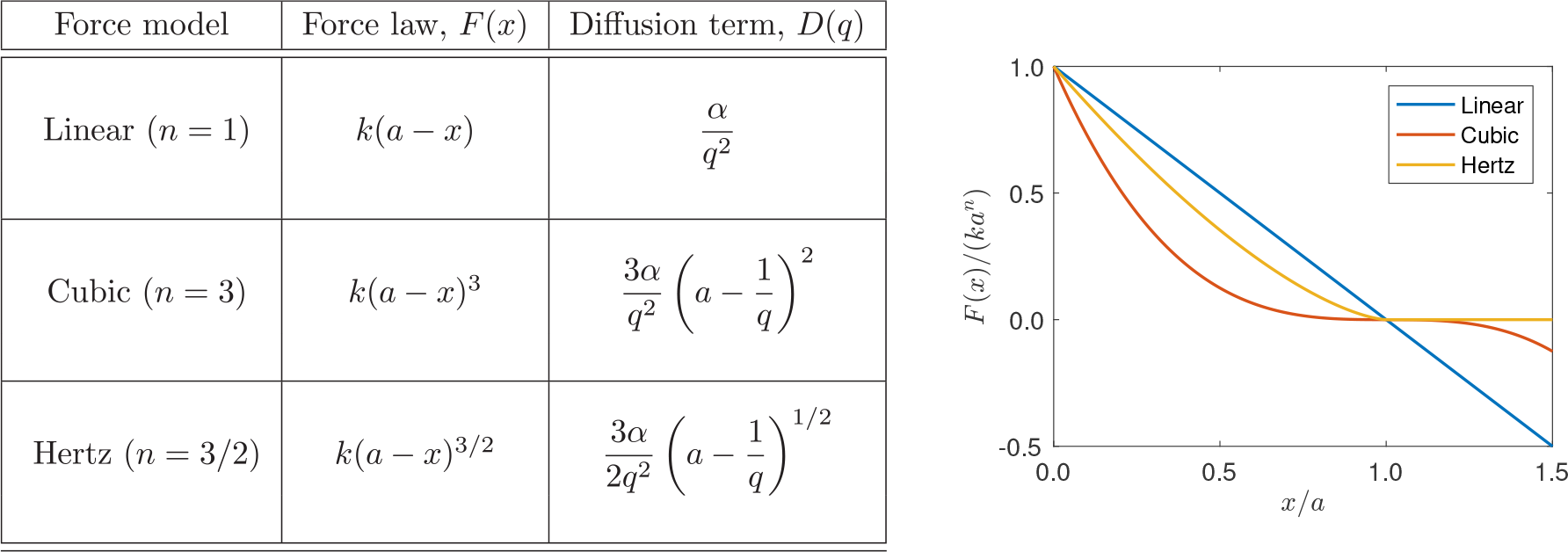
The force laws, *F*(*x*), and the corresponding diffusion coefficient, *D*(*q*), considered in Section 4.1.

## 4. General force law model

In this section we generalise the 1D cell-based model to account for a more general force law, *F*_*i*,*i*±1_, between neighbouring nodes, *i* and *i* ± 1, in equation (2)-(4). For concreteness, in our examples we will work with a force law of the form [15]

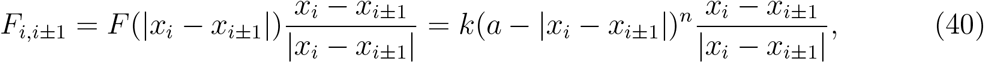

for some real valued exponent, *n*, where *n* = 1 gives a linear force law, as considered in Section 3, *n* = 3 gives a cubic force law, and *n* = 3/2 gives the Hertz force law (see Figure 3). These force laws are chosen to cover a wide range of potential cell interactions. The nodes evolve over time according to

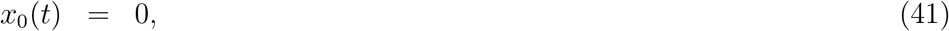

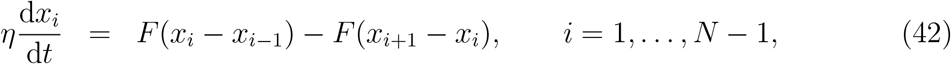

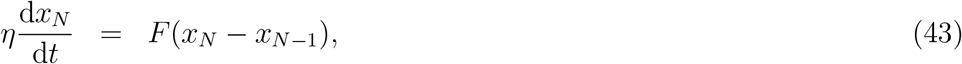

where *F*(*x*) = *k*(*a* − *x*)^*n*^. The initial conditions are 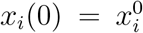 for *i* = 1,…, *N*. As for the linear force law case, we solve for the position of each node, *x*_*i*_, numerically using a simple forward Euler method with time-step Δ*t* = 0.001. Exemplar results for the model are shown in Figure 4, where we show how the leading edge and cell density of an initially compressed population of cells evolves over time for the three different force laws.

### 4.1. Continuum approximation

To derive a coarse-grained model we again, with a slight abuse of notation, extend node positions, *x*_*i*_(*t*), to a smooth function *x*(*i*, *t*), for *i* ∈ [0,*N*], and non-dimensionalise equations (41)-(43) using similar scalings to the simple linear case,

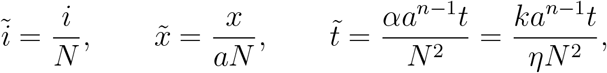

so that 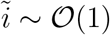 and 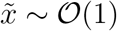. We also define the non-dimensional force function to be

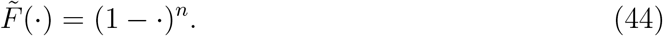

To derive an equivalent coarse-grained continuum model we proceed as in Section 3.1, performing a Taylor expansion about 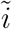 within the non-dimensionalised system to give, on neglecting terms which are 𝒪(1/*N*^2^),

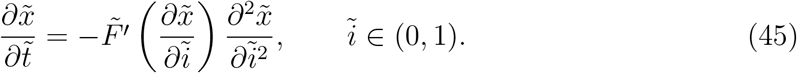

The left-hand boundary condition remains as 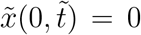 and, as before, to derive the right-hand boundary condition we Taylor expand and neglect terms which are 𝒪(1/*N*^2^) to give, at 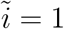,

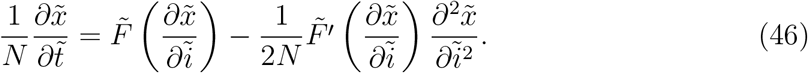

Rewriting in terms of dimensional variables we have the following PDE for *x*(*i*, *t*):

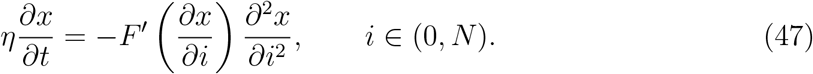

The boundary conditions are

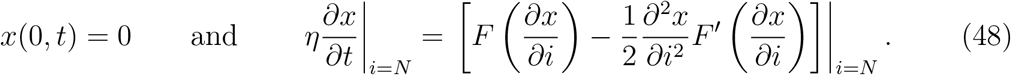

As before, the initial conditions can be specified by extending the discrete initial conditions, 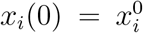 for *i* = 1,…, *N*, to a continuous function *x*(*i*, 0) such that 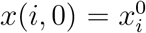 for *i* = 1,…, *N*. As a consistency check, we note that when *n* = 1, as for the linear force law, equations (47) and (48) reduce to equations (16) and (17).

### 4.2. Derivation of the corresponding cell density model

We can establish a PDE describing the evolution of cell density with position, x, and time, t, by making the same change of variables as in Section 3.2. Following a simple substitution of terms from equations (19) and (20) into equation (47) we obtain the PDE

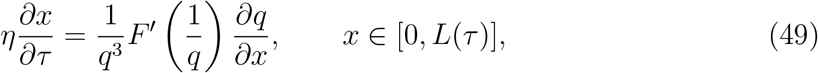

which represents how the domain, *x*, evolves along characteristics with constant index. Substitution of equations (19) and (20), and a simple rearrangement (multiplying by *q* and differentiating with respect to *x*, identical to earlier arguments), results in a PDE for the cell density of the form

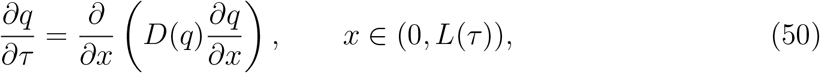

where the diffusion coefficient, *D*(*q*), is defined as

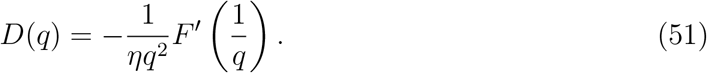

The characteristic equation (49) can then be rewritten as

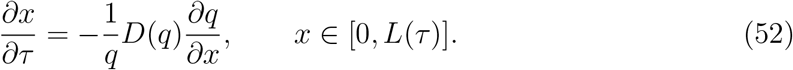

Under the same change of variables, boundary conditions become

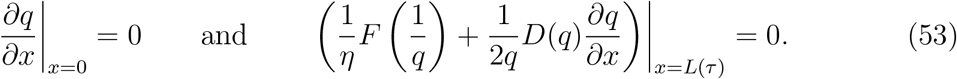

As in Section 3.2, we note that the system conserves total cell density:

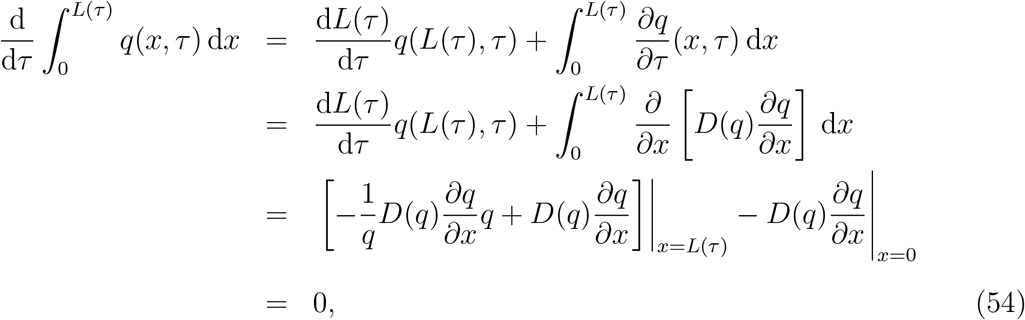

where the final result is established using equation (52).

#### 4.2.1. Numerical solution

In summary, the coarse-grained model consists of a PDE for the evolution of cell density,

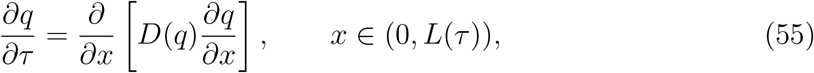

together with the boundary conditions

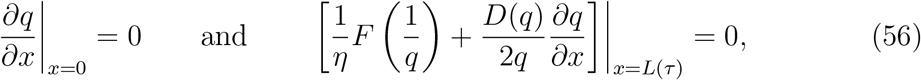

and initial condition

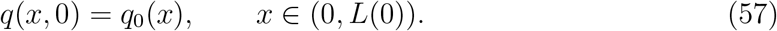

The characteristic equation is

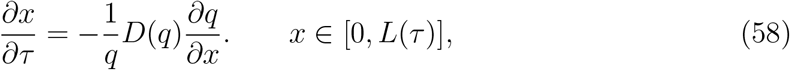

and we can use it to specify the evolution of the domain with time. In particular, we have

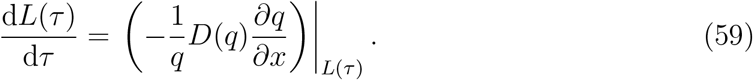

As in Section 3.2.1, in order to solve the coarse-grained model numerically, we employ a Lagrangian transformation to map the free boundary problem to a fixed domain: we let *τ* = *T* and

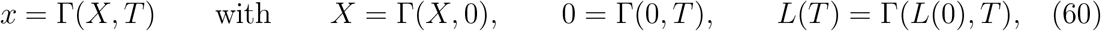

to give

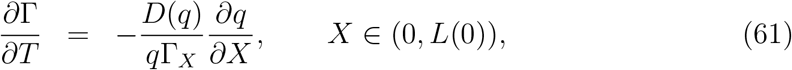

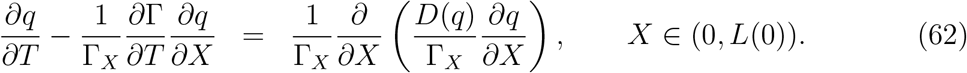

The initial and boundary conditions for Γ(*X*, *T*) are specified in equation (34), and for *q*(*X*, *T*) we have

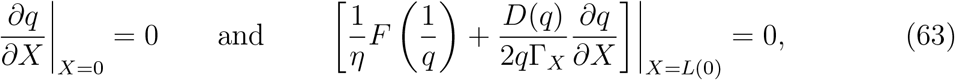

together with

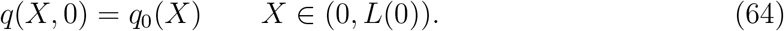

We solve the model numerically using an implicit finite difference method with Picard iteration. Full details are given in Appendix A.

### 4.3. Results

Across all force laws tested, the coarse-grained PDE model is very accurate in its prediction of both evolution of the cell density, *q*(*x*, *τ*), and the free boundary at *x* = *L*(*τ*) (see Figure 4). The only minor deviation in the predictions of the models is found at the leading edge, where the gradient in the cell density is largest. Note that in this region the approximation of the density in the cell-based model is lower order in *N*, so this deviation could perhaps be reasonably expected.

**Figure 4:**
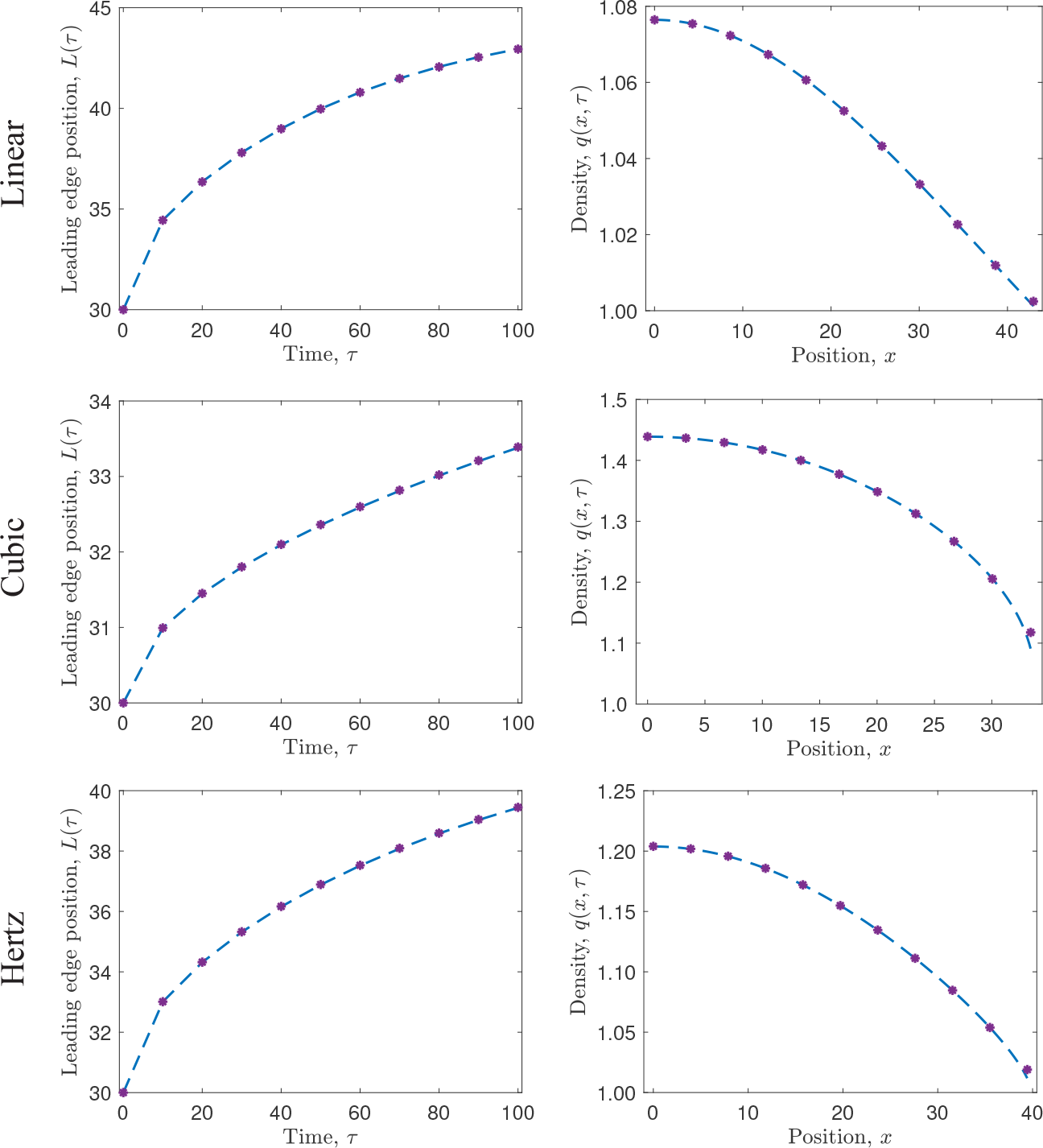
Comparison of the leading edge position, *L*(*τ*), and cell density, *q*(*x*, *τ*), predicted by the cell-based model, equation (41)-(43) with the force law as defined in (40) (purple asterisks), and the coarse-grained PDE model, equation (55)-(59) (blue dashed line), as the force law is varied (see Figure 3). In each case, *N* = 45 cells are initialised uniformly in *x* ∈ (0, 30), and *aN* = 45 and *α/a*^1−*n*^*N*^2^ = 135 are fixed.

## 5. Introducing proliferation into the model

In this section, we extend the model to include proliferation. The mechanism that we incorporate is stochastic, with each cell dividing with a defined rate per unit time. We derive a coarse-grained PDE model to describe the evolution of the domain, and cell density therein, over time, and demonstrate the validity of the coarse-grained model by comparing its solution to averaged results from the cell-based model.

### 5.1. Proliferation mechanism

To extend the model to include proliferation, we assume that each cell proliferates stochastically at a rate per unit time that is a function of its length. That is, the probability that cell *i* divides in the time interval [*t*, *t* + d*t*) is *G*_*i*_d*t* where *G*_*i*_ = *G*(|*x*_*i*_ − *x*_*i*−1_|). When a cell proliferates, a new node is introduced at its centre to establish the daughter cells, and we relabel the node indices to ensure their order, that is, *x*_*i*_(*t*) < *x*_*i*+1_(*t*) for *i* = 0,…, *N*(*t*) and *t* ≥ 0. Subsequently, when a new node (and cell) is introduced due to the proliferation of the *i*^th^ cell, we relabel the nodes with indices *j* = *i* + 1,…, *N* using *j* ↦ *j* + 1, as shown in Figure 5. In this work, we explore the dynamics introduced by three different types of proliferation mechanism: (i) cells proliferate at constant rate; (ii) cells proliferate at rate proportional to their length; and (iii) cells proliferate once they have reached a target length^3^. Specific functional forms for the growth rates we consider in this work are provided in Figure 6.

**Figure 5:**
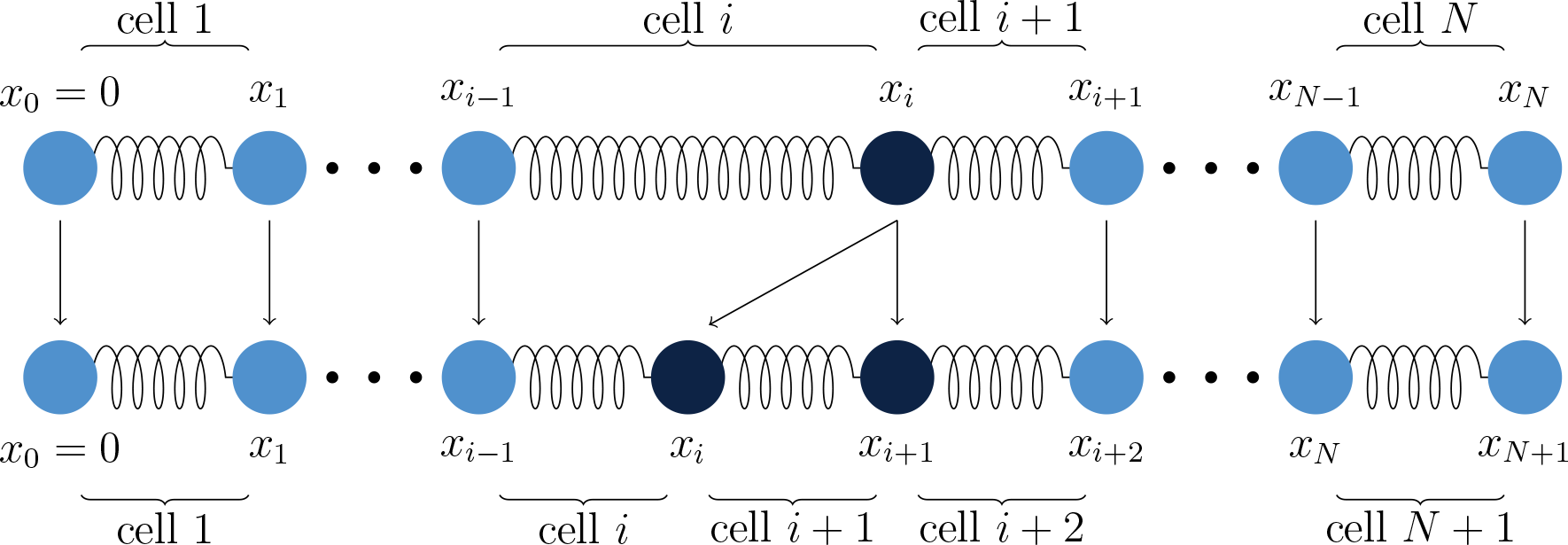
Proliferation of cell *i* entails the introduction of a new node at the cell centre, and relevant nodes and cells are then relabelled to ensure *x*_*j*_ < *x*_*j*+1_ for *j* ∈ {0,1,…, *N*}.

**Figure 6:**
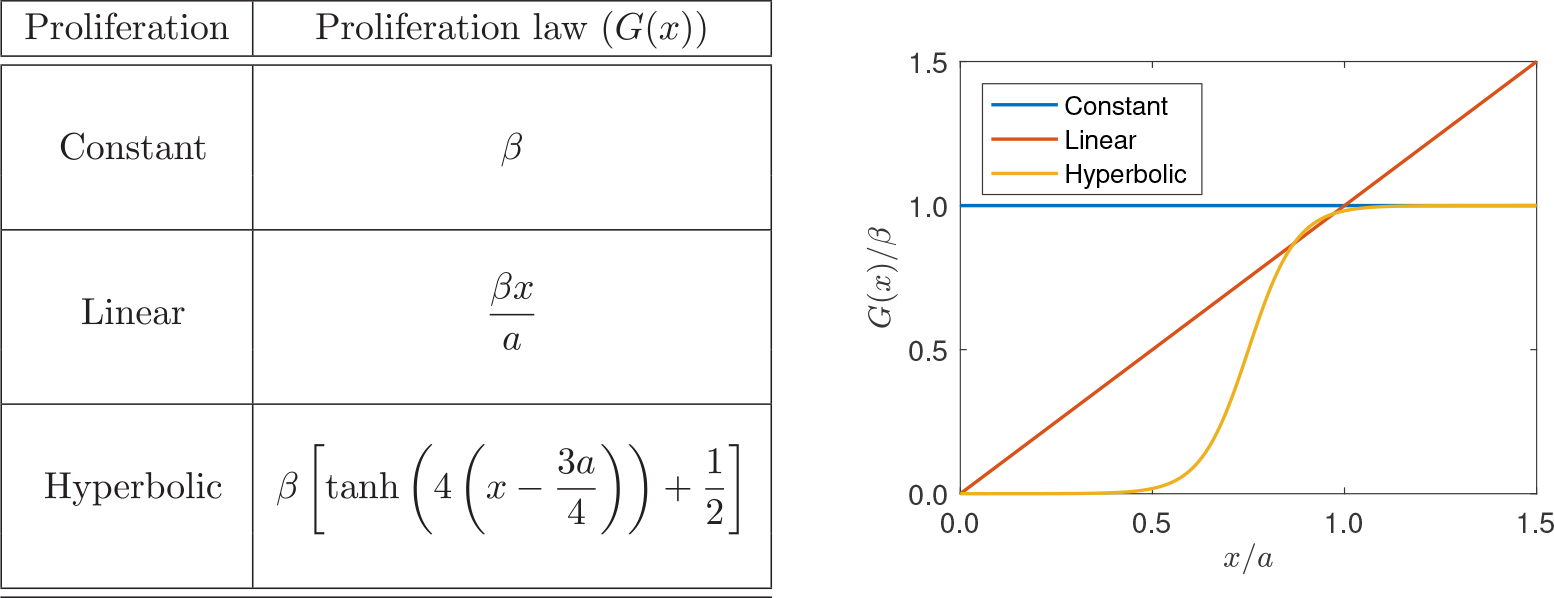
The proliferation laws considered in Section 5.

To generate individual realisations of this discrete stochastic model we use a constant time-step algorithm, with time-step Δ*t* = 0.001. At each step, we first update the position of each node, *x*_*i*_, *i* = 1,…, *N*, by using a simple forward Euler method to integrate equations (41)-(43) numerically, then we check to see whether a cell proliferation event occurs (and, if so, which cell proliferates). A cell proliferation event occurs with probability 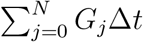 and, given a cell proliferation event occurs, the probability that cell *i* proliferates is 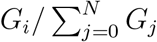, *i* = 1,…,*N*. In each case, we use rejection sampling to implement the decision, and if a cell proliferation occurs, we update the node indices as indicated in Figure 5. Note that this algorithm enforces the condition that at most one cell can proliferate per time-step; this is a reasonable approximation for the parameters used in this work.

### 5.2. Derivation of cell density model with proliferation

As the proliferation mechanism we have introduced is stochastic, we now consider evolution of the expected positions of the nodes over time. We make progress by considering the system over an infinitesimally small time interval [*t*, *t* + d*t*) and condition on cell proliferation taking place during that time interval to write, for *i* = 1,…, *N* − 1,

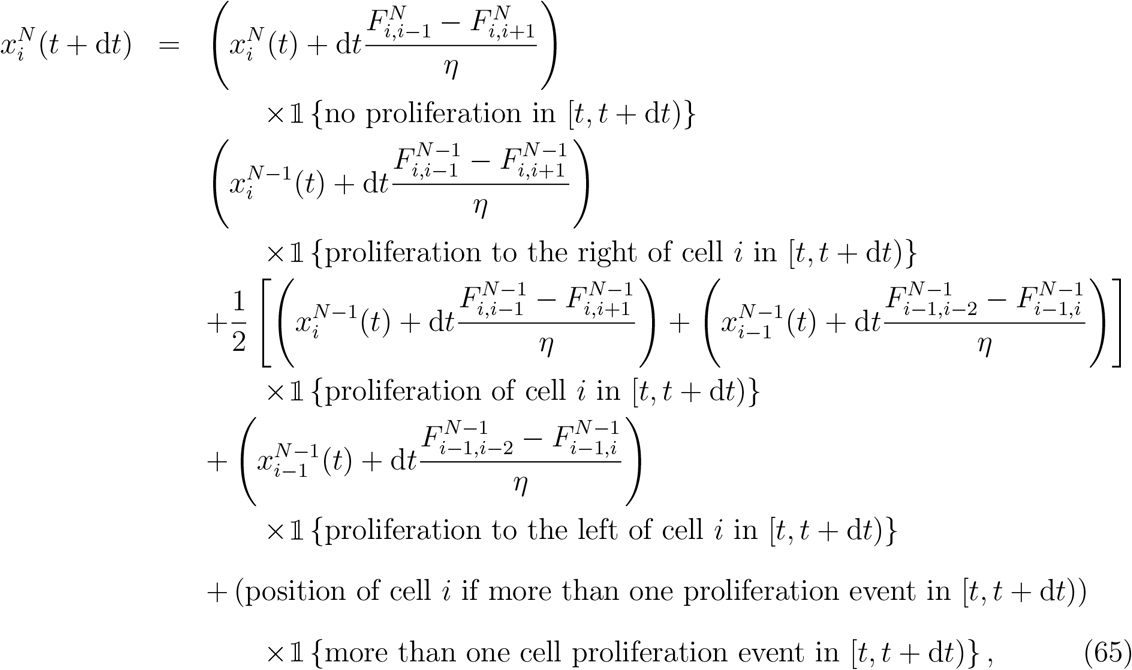

and

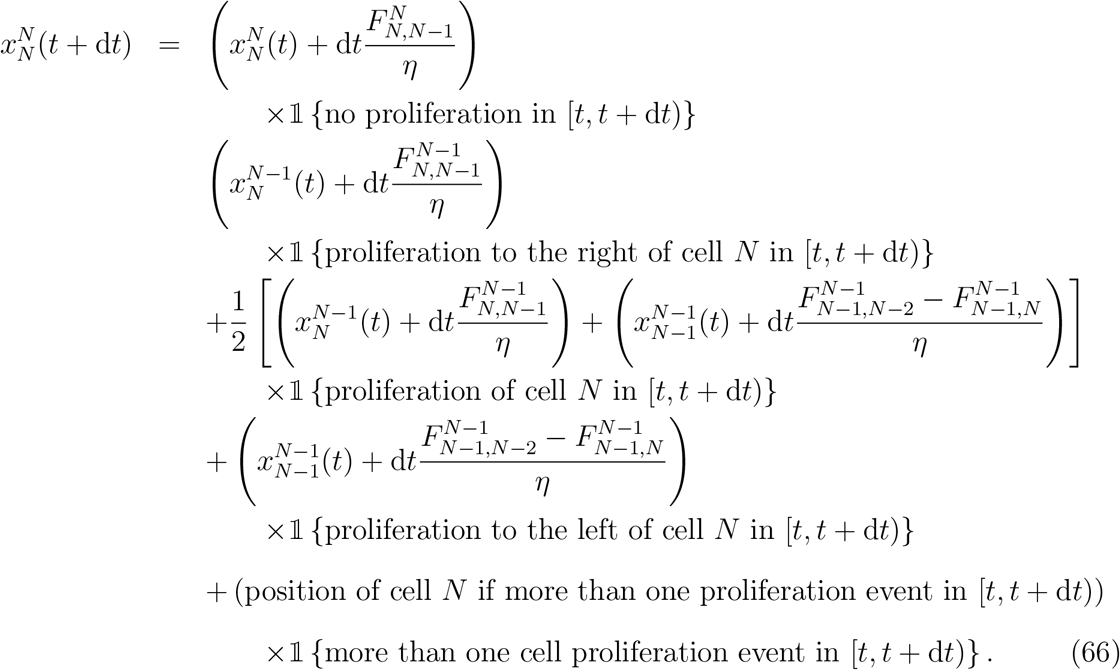

In equations (65) and (66) and 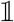 is the indicator function and 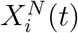 is the position of cell *i* at time *t* when there are *N* cells. The superscript *N* on the force terms, *F*_*i,j*_, indicate that they are evaluated using the positions of cells *i* and *j* when there are *N* cells.

The required probabilities to specify the indicator functions are

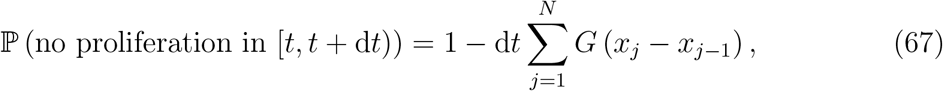

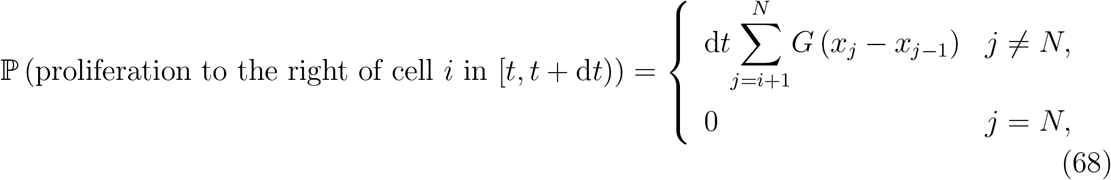

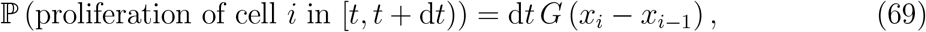

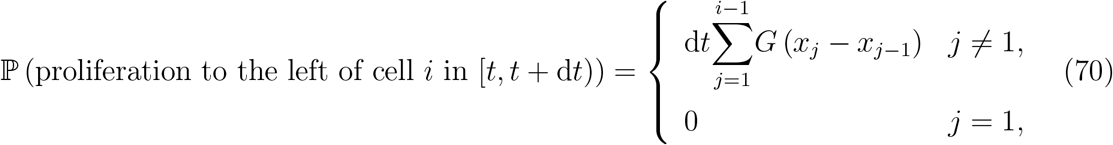

and

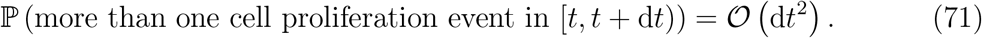

We now take expectations on both sides of equations (65) and (66), denoting by 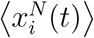 the expected position of node *i* at time *t* when there are *N* nodes. We then make two simplifying assumptions: (i) that 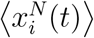 is a continuous function of time; and (ii) that 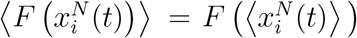 and 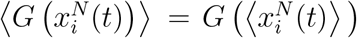. The former allows us to rearrange and take the limit as d*t* → 0 and, together with the latter, we have

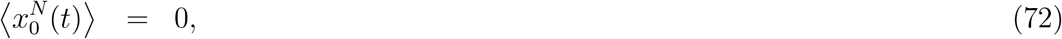

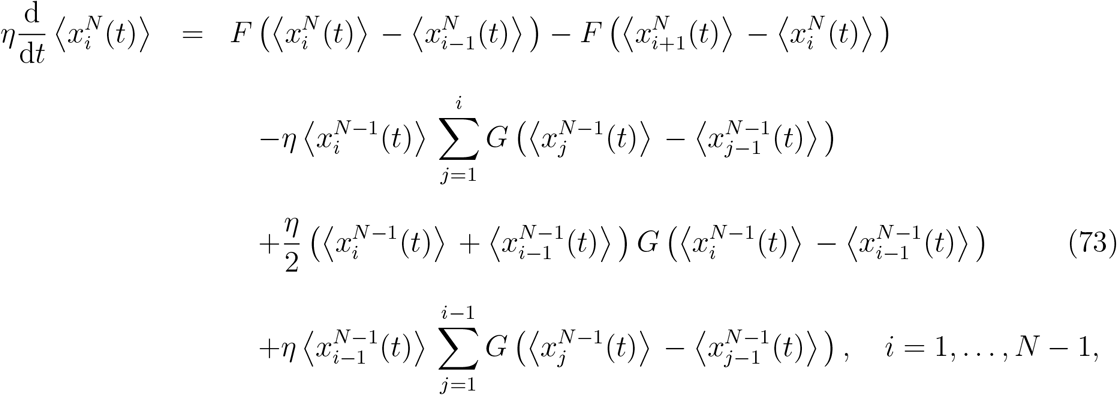

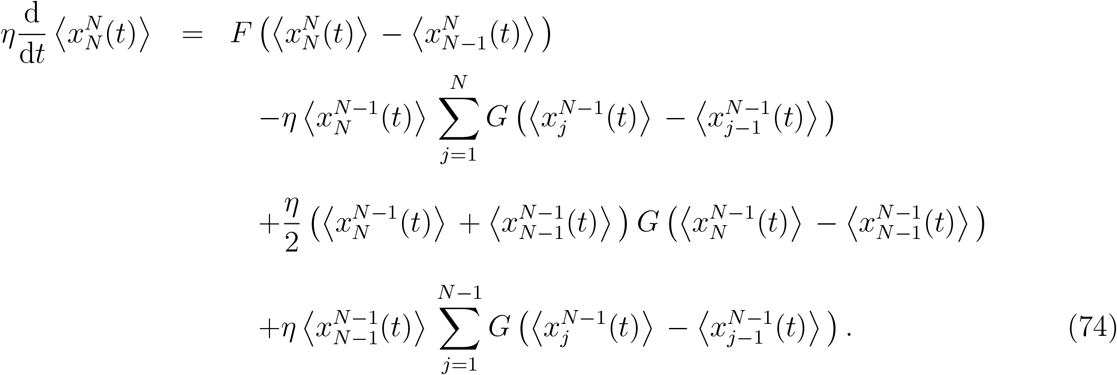

### 5.3. Continuum approximation

To enable a continuum approximation to be formulated we make the further approximation 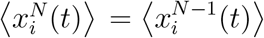, and to simplify exposition going forward, we will drop use of the angle brackets. After algebraic simplification, we have

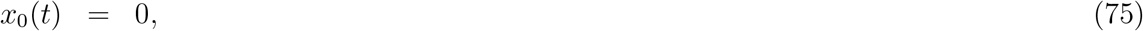

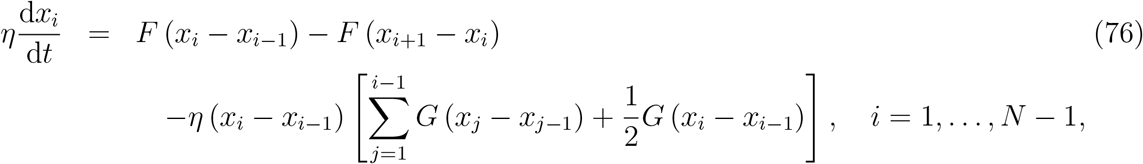

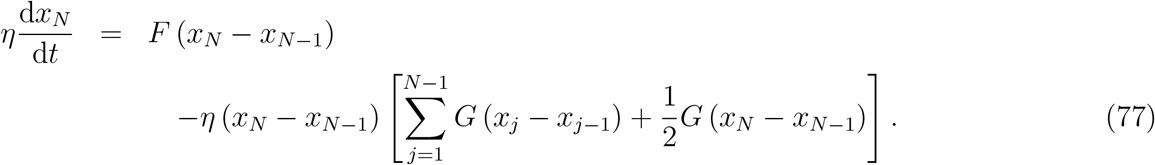

To make progress in deriving an equivalent continuum, coarse-grained model, we proceed as before: we extend node position, *x*_*i*_(*t*), to a smooth function *x*(*i*, *t*) for *i* ∈ [0, *N*(*t*)], and non-dimensionalise using the scalings

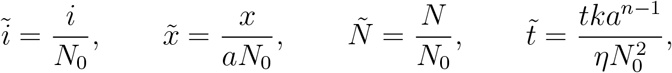

where *N*_0_ = *N*(0), the number of nodes at *t* = 0. We also define the non-dimensional proliferation function such that

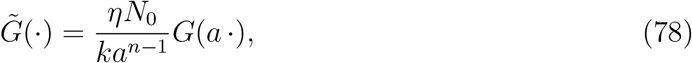

and the non-dimensional force function is as specified in equation (44).

We then work in the same manner as before, using Taylor expansion together with approximations of the form

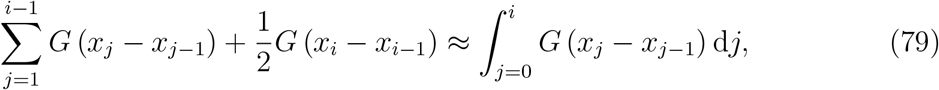

to give, upon neglecting terms that are 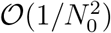,

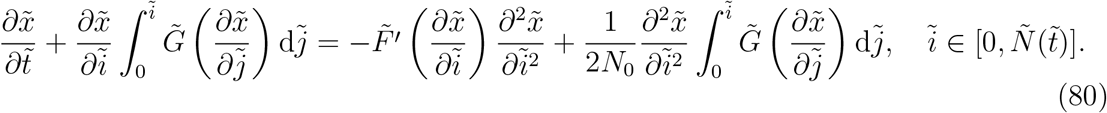

The left-hand boundary condition remains as 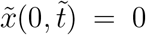 and, once again, we derive the right-hand boundary condition by Taylor expanding and neglecting terms that are 𝒪(1/*N*^2^) to give, at 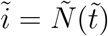,

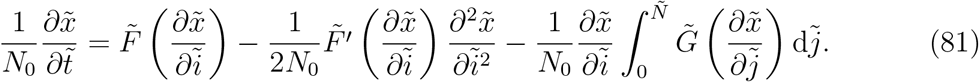

Rewriting equation (80) in terms of the dimensional variables we have the following PDE for *x*(*i*, *t*):

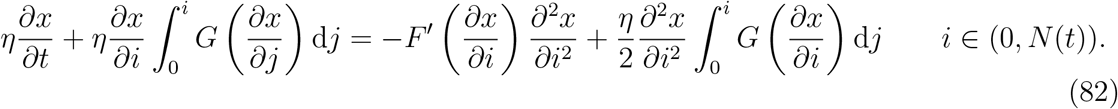

The boundary conditions are

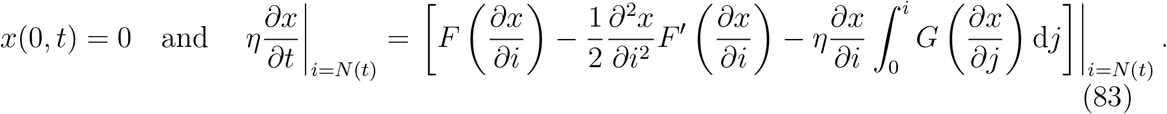

As before, the initial conditions can be specified by extending the discrete initial conditions, 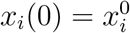 for *i* = 1,…, *N*(0), to a continuous function *x*(*i*, 0) such that 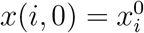 for *i* = 1,…, *N*(0).

### 5.4. Derivation of the corresponding cell density model

We now establish a PDE describing the evolution of cell density with position, *x*, and time, *t*, for a proliferative cell population with general force and proliferation laws. Changing variables from (*i*, *t*) to (*x*, *τ*), as before, with a simple substitution of terms from equations (19) and (20) into equation (82) we obtain the PDE

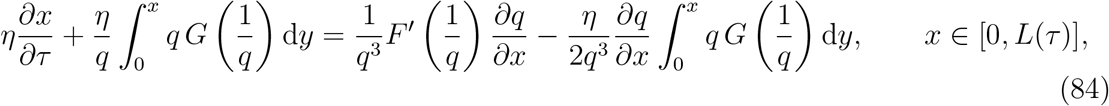

which represents how the domain evolves along the characteristics. Note that, due to proliferation, this is no longer equivalent to following constant cell index, *i*.

After further rearrangement, as before, we have

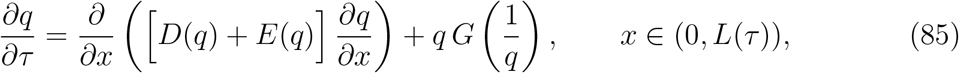

where

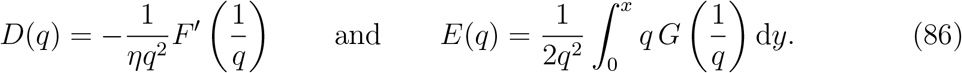

Under the same change of variables, the boundary conditions become

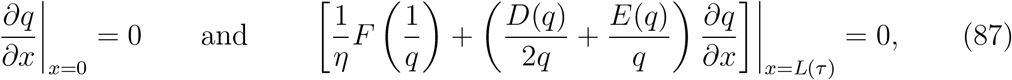

and the initial conditions, computed as in Section 3.1, are

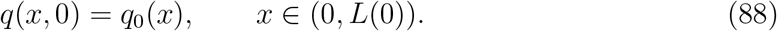

Using equation (86), characteristic equation (84) can be re-written as

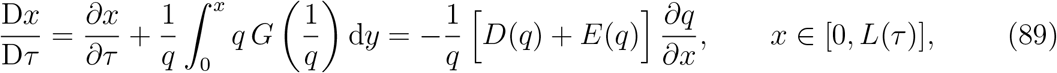

That the left-hand side of equation (89) constitutes a material derivative can be seen by following a small “tissue element” as the cell population grows and divides. We have *x* = *x*(*i*(*t*),*t*) with

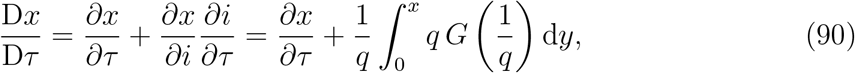

where we have used the fact that the rate of change of cell index is equal to the rate of cell proliferation in the region to the left of the cell *i.e.*

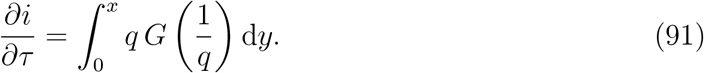

Finally, using equation (89) we can specify the rate of growth of the domain over time as

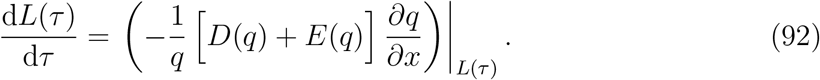

### 5.5. Evolution of cell number

Note that, since cell proliferation is now present in the model, the cell number changes over time and the system does not conserve mass. The cell number at time *τ* is specified by equation (18) as

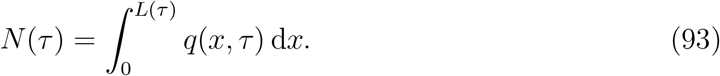

Differentiating with respect to *τ* and using the left-hand boundary condition (97) gives

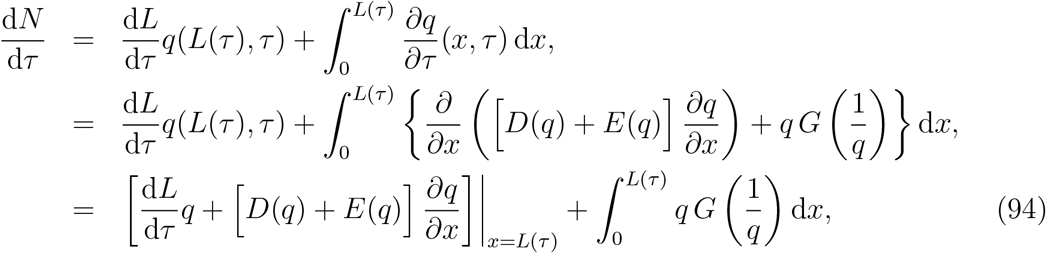

Substituting equation (92) into equation (94) gives the rate of change of cell number over time a

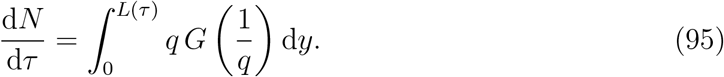

This equation states simply and intuitively that the rate of change of cell number is simply equal to the sum of the proliferation rates of each cell.

### 5.6. Numerical solution

In summary, the coarse-grained model consists of a PDE for the evolution of cell density

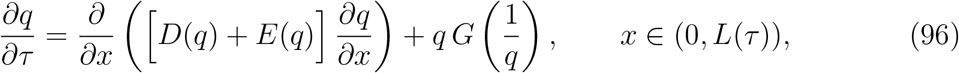

together with boundary conditions

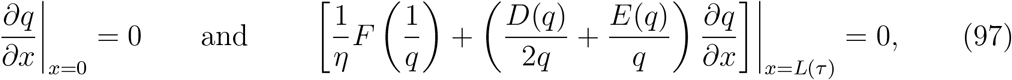

and initial condition

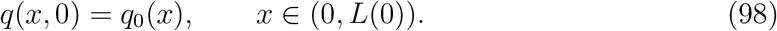

The characteristic equation is

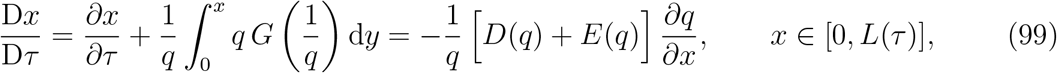

and we have

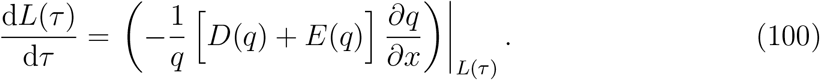

As in Section 3.2.1 and Section 4.2.1, in order to solve the coarse-grained model numerically, we employ a Lagrangian transformation to map the free boundary problem to a fixed domain: we let *τ* = *T* and

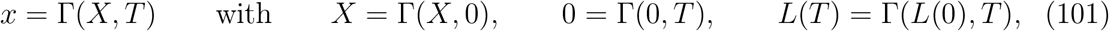

to give

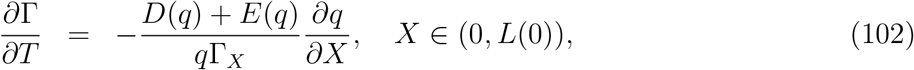

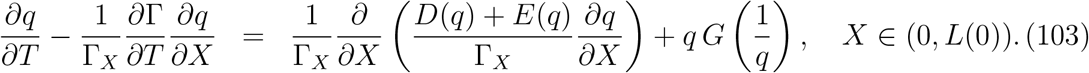

The initial and boundary conditions for Γ(*X*, *T*) are specified in equation (101), and for *q*(*X*, *T*) we have

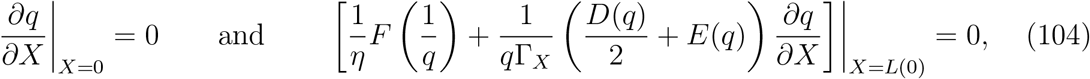

together with

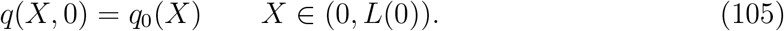

We solve the model numerically using an implicit finite difference method with Picard iteration. Full details are given in Appendix A.

### 5.7. Results

To demonstrate the validity of the coarse-grained model, we compare the solution of the PDE system, equation (96)-(100), with 100 averaged realisations of the discrete, stochastic model. Across all force laws and proliferation functions tested, the coarsegrained PDE model is very accurate in its prediction of both evolution of the mean cell number, *N*(*τ*), and the mean position of the free boundary at *x* = *L*(*τ*) (see Figure 7 and Figure 8, respectively)^4^. The different force laws and proliferation functions result in quite different behaviours, in particular how quickly the leading edge expands or how rapidly the number of cells grows.

## 6. Discussion and outlook

In this work we study a one-dimensional cell-based model of an epithelial sheet of cells where individual cells move deterministically and proliferate stochastically. This cell-based mechanical model gives rise to a moving boundary problem on the domain 0 < *x* < *L*(*τ*), where *τ* is time. We construct a continuum-limit description of the cell-based model, leading to a novel moving boundary PDE description governing the density of the cells within the evolving domain, 0 < *x* < *L*(*τ*), as well as a moving boundary condition governing the evolution of *L*(*τ*). Our results show that care must be taken to arrive at a moving boundary condition that conserves mass appropriately.

**Figure 7:**
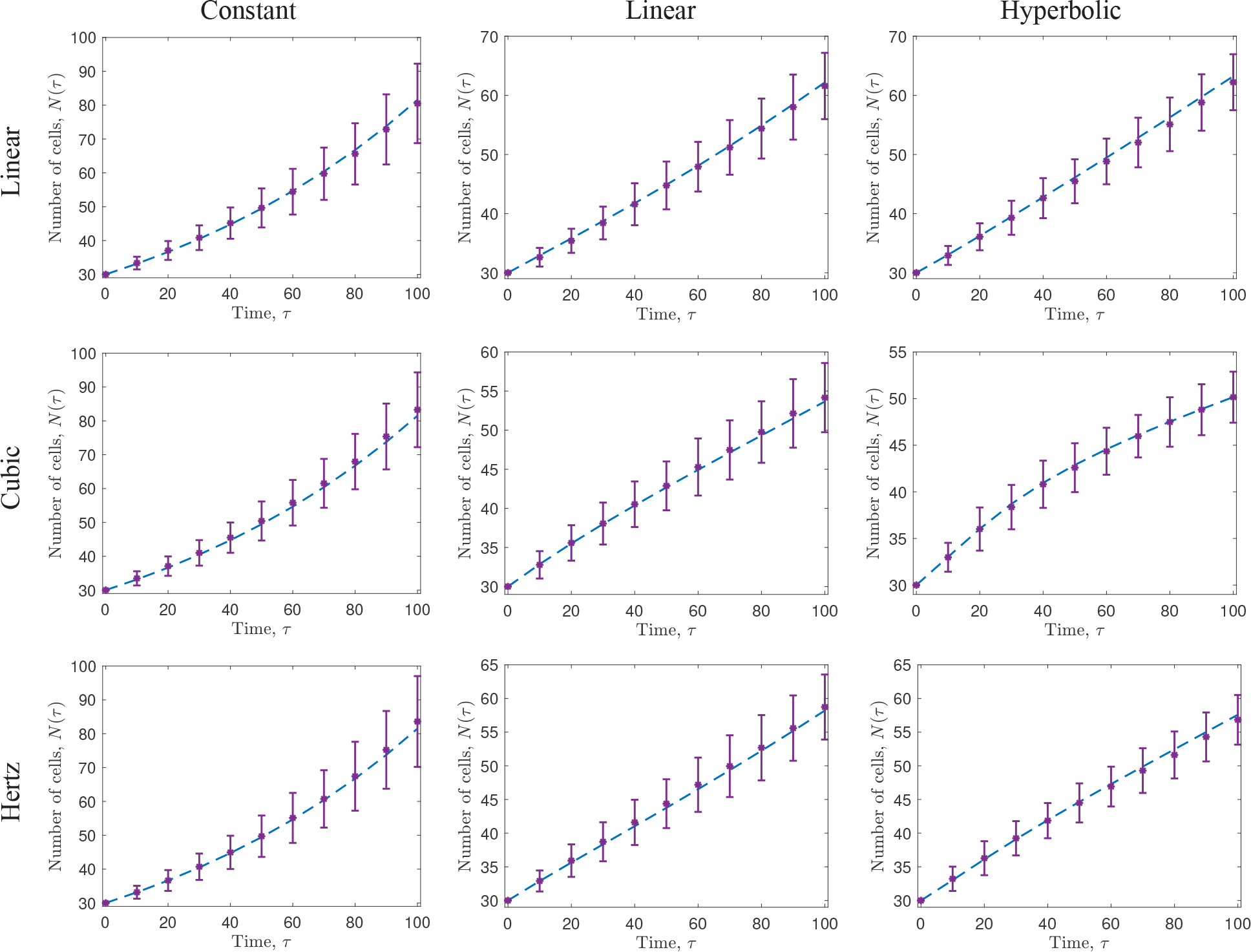
Comparison of the number of cells, *N*(*τ*), predicted by the cell-based model described in Section 5.1 (purple asterisks, and accompanying error bars), and the coarse-grained model, equation (96)-(100) (blue dashed line), for varying force laws and proliferation functions. Each row represents a different force law, whereas each column represents a different proliferation function. Each force law is defined and visualised in Figure 3, and each proliferation function is defined and visualised in Figure 6. In each case, we display averaged results from 100 realisations of the stochastic model, *N* = 30 cells are initialised uniformly in *x* ∈ (0, 30) and *a* =1, *α* = 15, and *β* = 0.001.

**Figure 8:**
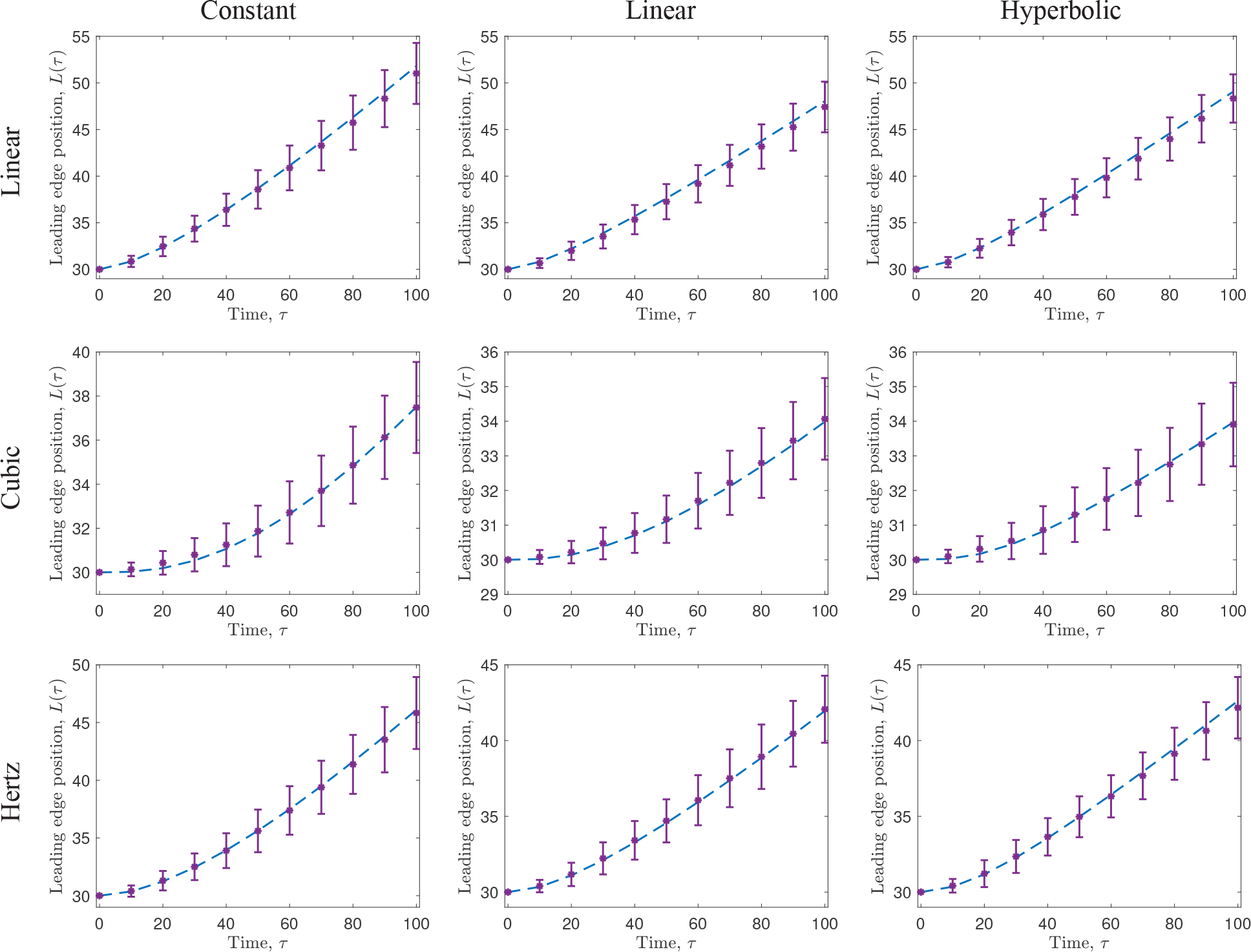
Comparison of the leading edge position, *L*(*τ*), predicted by the cell-based model described in Section 5.1 (purple asterisks, and accompanying error bars), and the coarse-grained model, equation (96)-(100) (blue dashed line), for varying force laws and proliferation functions. Each row represents a different force law, whereas each column represents a different proliferation function. Each force law is defined and visualised in Figure 3, and each proliferation function is defined and visualised in Figure 6. In each case, we display averaged results from 100 realisations of the stochastic model, *N* = 30 cells are initialised uniformly in *x* ∈ (0, 30) and *a* = 1, *α* =15, and *β* = 0.001.

There are many ways that our modelling approach can be extended, both from a theoretical point of view and a biological point of view. In all cases considered, we always study problems leading to an expanding population of cells where *L*(*τ*) is an increasing function of time. While these sets of problems are biologically relevant since they correspond to growing tissues, an interesting extension of our work would be to consider incorporating cell death and cell extrusion so that the model can be used to study both tissue growth and tissue shrinkage [25]. Other avenues for interesting extensions would be to consider the incorporation of internal boundaries within a mixed heterogeneous population so that the model could be used to study the interactions between an invasive population, such as a population of tumour cells, that invades into a surrounding population of non-invasive cells [6]. Furthermore, an obvious extension of the current work would be to two or three dimensions [22, 26]. In terms of biological applications, mechanical models describing cell migration and cell proliferation are important in wound healing [4], development [10], and cancer progression [20] and detection [7]. In all of these various applications we expect that experimental and clinical data will encompass both individual cell-based information as well as population-level, tissue-scale information. Therefore, the general framework of developing and applying cell-based models to study a particular phenomena while simultaneously working with a coarse-grained approximation to provide population-level information will be important to ensure that we get the most out of taking a combined modelling and experimental approach to studying particular biological phenomena.

## Acknowledgements

REB is a Royal Society Wolfson Research Merit Award holder and would like to thank the Leverhulme Trust for a Research Fellowship and and also acknowledges the BBSRC for funding via grant no. BB/R000816/1. AP would like to thank the UK’s Engineering and Physical Sciences Research Council (EPSRC, EP/G03706X/1) for funding through a studentship at the Systems Biology programme of The University of Oxford’s Doctoral Training Centre. MJS appreciates support from the Australian Research Council (DP170100474).

## Appendix A. Numerical solution of the full system

We solve equations (101)-(105) numerically using a finite difference scheme with Picard iteration. Algorithm 1 provides the scheme used for Picard iteration. The tolerance for the Picard iteration step is ϵ = 10^−4^, where the distance between solutions, *d*(*q*^*k*^,*q*^*k*+1^), is computed using the sum of squared differences over all space points, and the space step and time step are d*X* = 0.01 and d*τ* = 0.001, respectively.

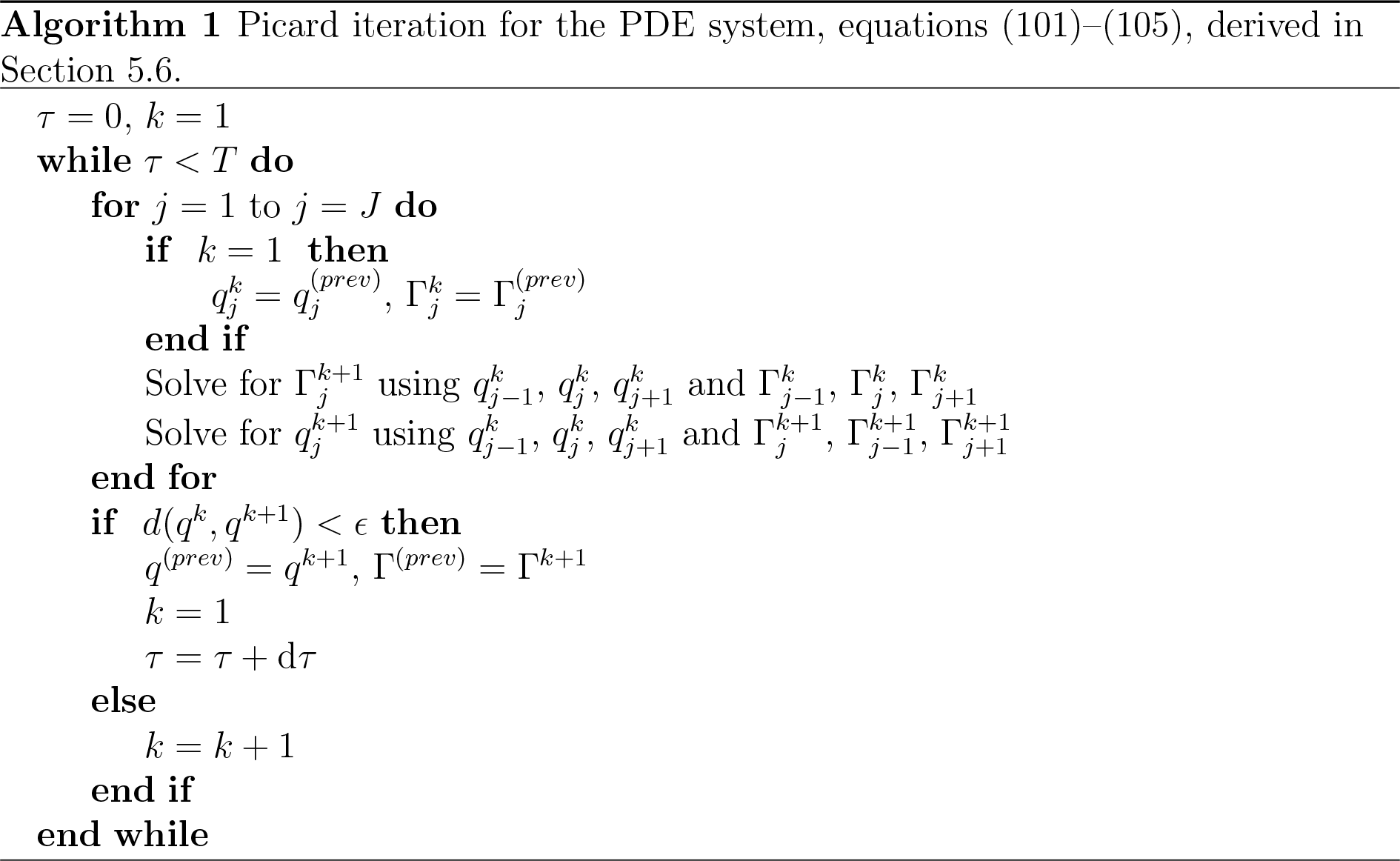

Within the Picard algorithm, equation (102) is discretised as

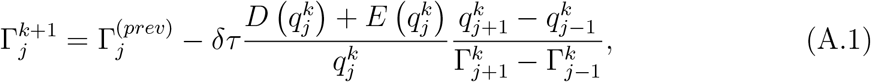

and equation (103) as

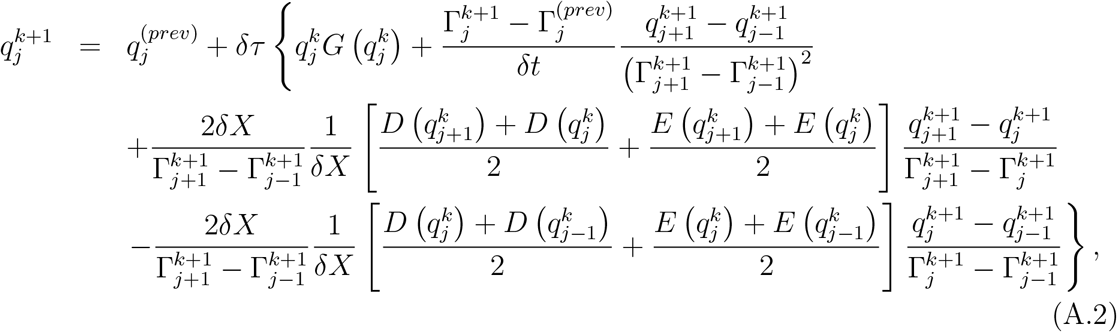

where the subscript, *j*, indexes the space step, the superscript, *k*, the Picard iteration step. Equation (A.2) can be rearranged to give

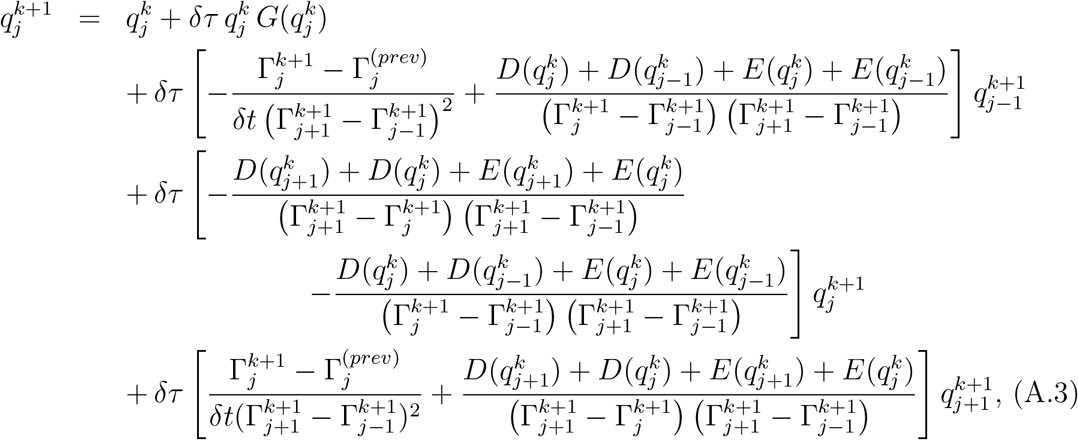

and then solved using the tridiagonal matrix algorithm.

## Footnotes

1 Throughout this work, we will assume 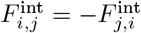.

2 Note that *x*(*i*, *t*) is a continuous variable representing position, has dimensions of length, and ranges between 0 and *N* × *a* when cells are at equilibrium.

3 Note that to avoid discontinuities, in this case we use a smoothed version of the Heaviside function, as given in Figure 6.

4 For example, the agreement of the models for varying *α*/*β* (doubled and halved) is also excellent (results not shown).

